# Wild edible yams from Madagascar: New insights into nutritional composition support their use for food security and conservation

**DOI:** 10.1101/2023.07.19.549652

**Authors:** Mirana K. Ratsimbazafy, Paul A. Sharp, Louisette Razanamparany, Mamy Tiana Rajaonah, Feno Rakotoarison, Kholoud K. Khoja, Paul Wilkin, Melanie-Jayne R. Howes

## Abstract

Yams (*Dioscorea* species) are an important food resource in Madagascar, where both cultivated winged yam (*D. alata*) and wild edible yams are consumed. However, there is limited knowledge on the nutrient composition of wild edible yams in Madagascar, and on how they compare with the cultivated winged yam. Therefore, in this study, nine wild edible yam species, one with two subspecies from Madagascar (*D. bako*, *D. buckleyana*, *D. irodensis*, *D. maciba*, *D. orangeana*, *D. pteropoda*, *D. sambiranensis* subsp. *bardotiae* and subsp. *sambiranensis*, *D. seriflora* and *Dioscorea* species Ovy valiha), were analysed for their nutrient composition, compared with cultivated *D. alata*. They include 6/6 of the most favoured wild edible yam species in Madagascar. We present new nutrient composition data (protein, carbohydrate/starch, energy, lipid, β-carotene, minerals) for these nine wild edible yam species and show that they contain comparable levels of lipids and starch to *D. alata*, but none are better sources of protein than *D. alata*. We show that *D. irodensis* contains a significantly higher β-carotene content when compared to all other edible yams analysed, and that *D. buckleyana*, *D. irodensis* and *D. sambiranensis* subsp. *bardotiae* have a higher calcium content than cultivated *D. alata*, while all nine wild edible yam species analysed contain a higher iron content, compared to cultivated *D. alata*. The nutrient composition data presented could provide new incentives to conserve wild edible yams and inform on strategies to select *Disocorea* species for sustainable cultivation and use, providing opportunities to enhance future food security in Madagascar.

## 1. Introduction

Yam tubers, which are the underground organs of species in the plant genus *Dioscorea* (Dioscoreaceae), are important food resources in Madagascar, an island in the Indian Ocean located off the eastern coast of Africa. Thirty-eight *Dioscorea* species have been identified in Madagascar, representing 6% of the world’s diversity of this genus; thirty of these species are endemic and occur in the wild, and they are virtually all edible (Wilkin et al., 2022). Wild edible *Dioscorea* species are found in all parts of the island, both in humid areas such as in the east of the country, and in the more arid regions in the west and south.

While cultivated varieties of non-endemic *D. alata* L. (which is of south-east Asian origin) are available in Madagascar, their supply is limited; thus, rural communities harvest or buy locally sourced wild edible *Dioscorea* species as a supplementary food, or as a substitute for rice during periods of hardship and famine. Indeed, wild edible yams in Madagascar are an important seasonal food in certain regions such as Diana and Menable and are widely used in much of the island nation in times of hardship. Although yam tubers are an important food resource in Madagascar, there remains limited published data on the nutrient composition of wild edible yams endemic to Madagascar (Jeannoda *et al*. 2007). Of the *Dioscorea* species known to occur in Madagascar, those that have been evaluated have almost all been sourced from other countries, so their nutrient composition may not reflect that of tubers from species harvested in Madagascar. *D. alata* tubers from south Asia contain a high proportion of starch (65 - 84%) and approximately 7% protein (dry weight), although different cultivars may vary in their content of these macronutrients (Wanasundera & Ravindran, 1994; Rinaldo, 2020).

Wild edible yams in Madagascar are threatened by habitat degradation and by over or unsustainable use, thus risking food security and biodiversity loss, with some species (*D. bako*, *D. buckleyana*, *D. irodensis*, *D. orangeana*) already categorised as Endangered according to the International Union for the Conservation of Nature (IUCN) Red List (IUCN, 2022). *D. sambiranensis* subsp. *bardotiae* is also categorised as Endangered, while *D. sambiranensis* subsp. *sambiranensis* is categorised as Vulnerable (IUCN, 2022). *D. sambiranensis* is divided into two subspecies based on the vegetative and infructescence traits (Wilkin et al., 2009). These challenges to biodiversity conservation and to the role of wild edible yams in food security are exacerbated by the impact of climate change, which may result in increased temperatures, altered precipitation patterns leading to floods and drought and extreme weather events, which are already concerns in Madagascar, impacting on agriculture and the economy and food security (WHO and UN, 2021). The impact of climate change on food security is an additional concern that may intensify the triple burden of malnutrition (undernutrition, micronutrient deficiencies and obesity) and metabolic and lifestyle factors for diet-related non-communicable diseases; indeed, in 2021, 43.1% of the adult population in Madagascar were considered undernourished, while 4.5% of the adult population were considered obese, and 3.9% had diabetes (WHO and UN, 2021).

New knowledge of the nutritional value of wild edible yams in Madagascar may facilitate conservation actions by providing a scientific evidence base, and therefore incentives, for their sustainable cultivation and use, in addition to providing new insights into their nutritional value that could enable informed dietary choices. In this context, the aims of this study are to assess the nutrient composition of tubers from nine wild edible yam species, one with two subspecies, from Madagascar (*D. bako*, *D. buckleyana*, *D. irodensis*, *D. maciba*, *D. orangeana*, *D. pteropoda*, *D. sambiranensis* subsp. *bardotiae* and subsp. *sambiranensis*, *D. seriflora* and *Dioscorea* species Ovy valiha, an undescribed species), and to compare their nutrient content with that of the cultivated non-endemic winged yam, *D. alata*. The wild edible yams analysed include 6/6 (*D. bako*, *D. buckleyana*, *D. maciba*, *D. orangeana*, *D. sambiranensis*, *D. seriflora*) of the most favoured wild edible yam species in Madagascar (Wilkin et al., 2022).

## 2. Materials and methods

### 2.1 Samples

The tubers of nine wild edible yam (*Dioscorea*) species, in addition to cultivated *D. alata*, were harvested in Madagascar. The *Dioscorea* species collected, their geographical collection locations, common names and conservation status are shown in Table 1 (additional details, including GPS co-ordinates for collection locations and month and year of harvest, are shown in Supplementary Table 1). Harvested tubers were air-dried and sliced prior to analysis for nutritional composition. For nutrient analyses, samples were ground to a fine powder using a coffee grinder.

**Table 1.** Wild edible yam tubers collected: species, common names and collection locations†.

| *Dioscorea* species | IUCN Conservation Status<sup>‡</sup> | Common Name | Collection Location | District (Collection Location) |
| --- | --- | --- | --- | --- |
| *Dioscorea* species | Unknown | Ovy valiha | Andranomadiro | ANTSIRANANA |
| *D. sambiranensis* subspecies *bardotiae* Wilkin | Endangered | Ovy jaune | Betahitra |  |
| *D. pteropoda* Boivin ex H.Perrier | Vulnerable | Ovindambo | Ambilo |  |
| *D. buckleyana* Wilkin | Endangered | Taravy | Betahitra |  |
| *D. orangeana* Wilkin | Endangered | Ovy jia | Ivovona |  |
| *D. irodensis* Wilkin, Rajaonah & Randriamb. | Endangered | Ovy laglasy | Ivovona |  |
| *D. maciba* Jum. & H.Perrier | Least Concern | Ovy cable | Mahagaga |  |
| *D. sambiranensis* subspecies *sambiranensis* | Vulnerable | Angona | Ankariha | AMBANJA |
| *D. seriflora* Jum. & H.Perrier | Least Concern | Ovy sofy | Anaboranosalama |  |
| *D. bako* Wilkin | Endangered | Bako | Tsaraotana | MORONDAVA |

<sup>†</sup>Additional collection details are presented in Supplementary Table 1. <sup>‡</sup>IUCN (2022).

### 2.2 Protein analysis

Protein content was determined from the total nitrogen content by the Kjeldahl method (Bradstreet, 1965). In brief, 0.25g of powdered tuber sample was placed in a flask containing a solution of a concentrated sulfuric acid and Kjeldahl catalyst. After approximately 3 hours, when the solution was translucent, 50 ml of sodium hydroxide (40%) was added to neutralise the pH. After the distillation phase, the distillate was collected in aqueous boric acid solution (4%) prior to titration with sulfuric acid (0.1N). Protein content was estimated from the total nitrogen content of food using 6.25 as the conversion factor. Protein content was also determined using the colourimetric Bradford assay, as described previously (Maehre et al., 2018). Protein was extracted from each sample (500mg yam powder) using an extraction buffer (0.1M NaOH in 3.5% NaCl). The solution was vortexed and incubated for 30 minutes at 60°C, followed by centrifugation for 10min at 5000g. The supernatant was collected and protein content was measured with a commercially available assay kit (Bio-Rad, Inc.) using bovine serum albumin as a standard. Data are calculated as mg/g dry weight and presented as % values.

### 2.3 Carbohydrate and starch analysis

Total carbohydrate content was determined by subtracting the protein, lipid, ash and water content in the total weight of the tuber samples (FAO, 1998). Total starch content was measured using a Total Starch HK Assay Kit [K-TSHK] (Megazyme, Bray, Ireland). The assay followed the AOAC Official Method 996.11 for total starch determination (Englyst & Cummings, 1988; McCleary et al., 1994; McCleary et al., 1997). Samples were assayed in triplicate. Total starch content was calculated in accordance with the manufacturer’s (Megazyme, Bray, Ireland) instructions and data are presented as g/100g dry weight.

### 2.4 Energy analysis

The energy value of tuber samples was determined using the Atwater method, based on the heat combustion of nutrients (1g of protein provides 4 kcal / 17 kJ; 1g of fat provides 9 kcal / 37 kJ; 1g of carbohydrate provides 4 kcal / 17 kJ) (Atwater and Woods, 1896). The energy was also evaluated using the ballistic bomb calorimeter method to measure the gross energy content of each yam sample (Miller & Payne, 1959). Analysis was carried out using a Gallenkamp Ballistic Bomb Calorimeter [IKA©C1 Compact Calorimeter] and gross energy content is presented as kcal/100g dry weight.

### 2.5 Lipid analysis

Lipid content was determined using the Folch method (Folch *et al.,*1957). In brief, 1.0g tuber sample was added to a solution of chloroform/methanol (2/1, v/v) prior to homogenisation and vacuum filtration. The lower phase of the filtrate was recovered prior to evaporation at 65°C, followed by steaming at 85°C.

### 2.6 Ash analysis

Ash content was determined by placing 5.0g tuber material in a porcelain crucible prior to incineration in a muffle furnace at 550°C until white/grey ash consisting of inorganic material was obtained. After cooling in a desiccator, the mass of the ash was determined (AOAC International, 2000).

### 2.7 β-Carotene analysis

Carotenoids were extracted from samples of yam (500mg) using a ternary solvent system added, which consisted of 10ml hexane, 5ml methanol and 5ml HPLC water (Bóna-Lovász et al., 2013). The sample was first dissolved in hexane and methanol, followed by a 2-min vortex. Tubes were wrapped with foil in order to protect the carotenoids from daylight and incubated overnight. Subsequently, HPLC-grade water was added to each tube to separate the methanol from the solvent. The solution was then fully vortexed for 1 minute and centrifuged (3000rpm, 30 minutes) to further separate the hexane layer (containing the carotenoids). Following centrifugation, the upper hexane layer was collected and filtered. Absorbance was measured at 490 nm, and carotenoid content was measured using a standard curve with β-carotene as the reference compound. The data are expressed as µg β-carotene equivalents/100g dry weight.

### 2.8 Mineral analysis

Mineral analysis to quantify zinc, iron and calcium was performed using inductively coupled plasma-optical emission spectrometry (ICP-OES, Thermo-Fisher Scientific). Yam samples (50 mg) were mixed with 10 ml nitric acid (40%) and subjected to microwave digestion (CEM corporation, UK). The total volume of each tube was adjusted to 14 ml with HPLC-grade water and Yttrium (1 µg/ml final concentration) was added to each tube as an internal standard. Tubes were mixed and centrifuged for 5 min at 2000 rpm before analysis. Mineral content was calculated using standard curves generated using ICP-grade standards for iron, zinc and calcium and data are presented as mg/100g dry weight.

### 2.9 Statistical analysis

Data are presented as the mean ± SEM from at least three independent experiments. Statistical differences were determined using 1-way ANOVA followed by either Tukey’s (for multiple comparisons) or Dunnett’s (for comparison of wild yam species with *D. alata*) post-hoc tests where appropriate. Differences where P<0.05 were considered statistically significant. Statistical analysis was performed using SigmaPlot (version 14.5).

## 3. Results

### 3.1 Protein content

Nine wild edible *Dioscorea* species and cultivated *D. alata* from Madagascar were analysed over 12 harvest seasons from 2018 to 2020 (Table 2). Differences in protein content occurred between different harvest seasons, but protein content was not consistently high or low for any harvest season, thus indicating that harvest season is not a determinant for protein content in the *Dioscorea* species analysed. The mean protein content of cultivated *D. alata* (6.4g/100g dry weight [DW] 2018-2019) did not differ significantly from the mean protein content of any of the wild edible species analysed. In tubers harvested in 2019-2020, the mean protein content of *D. alata* (8.5g/100g DW) was significantly higher compared to all wild edible species analysed (*D. buckleyana* P<0.03; all other species P<0.01), except for *D. irodensis; D. sambiranensis* subsp. *bardotiae* and *D. bako*.

**Table 2.**
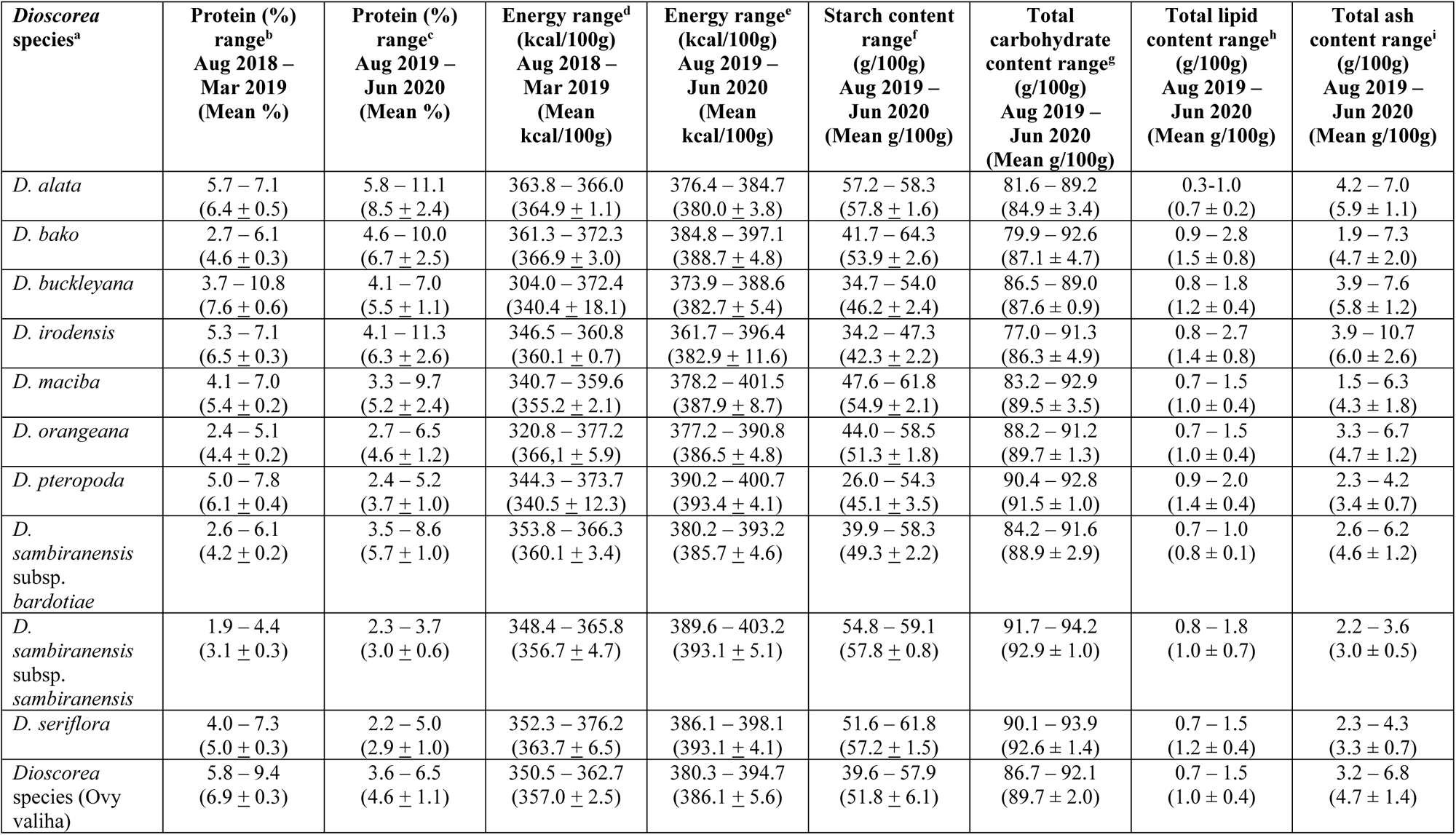
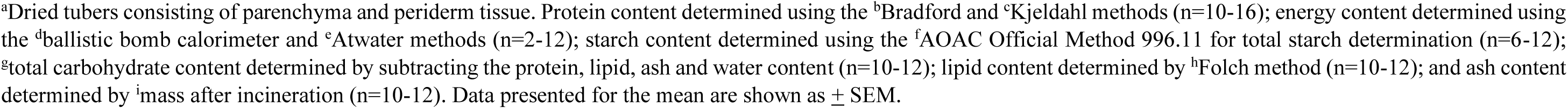
Protein, energy, starch, total carbohydrate, lipid and ash content of wild edible yams (*Disocorea* species) and cultivated winged yam (*D. alata*) collected over different harvest seasons in Madagascar, calculated as dry weight.

| *Dioscorea*<br>species<sup>a</sup> | Protein (%)<br>range<sup>b</sup><br>Aug 2018 –<br>Mar 2019<br>(Mean %) | Protein (%)<br>range<sup>c</sup><br>Aug 2019 –<br>Jun 2020<br>(Mean %) | Energy range<sup>d</sup><br>(kcal/100g)<br>Aug 2018 –<br>Mar 2019<br>(Mean<br>kcal/100g) | Energy range<sup>e</sup><br>(kcal/100g)<br>Aug 2019 –<br>Jun 2020<br>(Mean<br>kcal/100g) | Starch content<br>range<sup>f</sup><br>(g/100g)<br>Aug 2019 –<br>Jun 2020<br>(Mean g/100g) | Total<br>carbohydrate<br>content range<sup>g</sup><br>(g/100g)<br>Aug 2019 –<br>Jun 2020<br>(Mean g/100g) | Total lipid<br>content range<sup>h</sup><br>(g/100g)<br>Aug 2019 –<br>Jun 2020<br>(Mean g/100g) | Total ash<br>content range<sup>i</sup><br>(g/100g)<br>Aug 2019 –<br>Jun 2020<br>(Mean g/100g) |
| --- | --- | --- | --- | --- | --- | --- | --- | --- |
| *D. alata* | 5.7 – 7.1<br>(6.4 ± 0.5) | 5.8 – 11.1<br>(8.5 ± 2.4) | 363.8 – 366.0<br>(364.9 ± 1.1) | 376.4 – 384.7<br>(380.0 ± 3.8) | 57.2 – 58.3<br>(57.8 ± 1.6) | 81.6 – 89.2<br>(84.9 ± 3.4) | 0.3–1.0<br>(0.7 ± 0.2) | 4.2 – 7.0<br>(5.9 ± 1.1) |
| *D. bako* | 2.7 – 6.1<br>(4.6 ± 0.3) | 4.6 – 10.0<br>(6.7 ± 2.5) | 361.3 – 372.3<br>(366.9 ± 3.0) | 384.8 – 397.1<br>(388.7 ± 4.8) | 41.7 – 64.3<br>(53.9 ± 2.6) | 79.9 – 92.6<br>(87.1 ± 4.7) | 0.9 – 2.8<br>(1.5 ± 0.8) | 1.9 – 7.3<br>(4.7 ± 2.0) |
| *D. buckleyana* | 3.7 – 10.8<br>(7.6 ± 0.6) | 4.1 – 7.0<br>(5.5 ± 1.1) | 304.0 – 372.4<br>(340.4 ± 18.1) | 373.9 – 388.6<br>(382.7 ± 5.4) | 34.7 – 54.0<br>(46.2 ± 2.4) | 86.5 – 89.0<br>(87.6 ± 0.9) | 0.8 – 1.8<br>(1.2 ± 0.4) | 3.9 – 7.6<br>(5.8 ± 1.2) |
| *D. irodensis* | 5.3 – 7.1<br>(6.5 ± 0.3) | 4.1 – 11.3<br>(6.3 ± 2.6) | 346.5 – 360.8<br>(360.1 ± 0.7) | 361.7 – 396.4<br>(382.9 ± 11.6) | 34.2 – 47.3<br>(42.3 ± 2.2) | 77.0 – 91.3<br>(86.3 ± 4.9) | 0.8 – 2.7<br>(1.4 ± 0.8) | 3.9 – 10.7<br>(6.0 ± 2.6) |
| *D. maciba* | 4.1 – 7.0<br>(5.4 ± 0.2) | 3.3 – 9.7<br>(5.2 ± 2.4) | 340.7 – 359.6<br>(355.2 ± 2.1) | 378.2 – 401.5<br>(387.9 ± 8.7) | 47.6 – 61.8<br>(54.9 ± 2.1) | 83.2 – 92.9<br>(89.5 ± 3.5) | 0.7 – 1.5<br>(1.0 ± 0.4) | 1.5 – 6.3<br>(4.3 ± 1.8) |
| *D. orangeana* | 2.4 – 5.1<br>(4.4 ± 0.2) | 2.7 – 6.5<br>(4.6 ± 1.2) | 320.8 – 377.2<br>(366.1 ± 5.9) | 377.2 – 390.8<br>(386.5 ± 4.8) | 44.0 – 58.5<br>(51.3 ± 1.8) | 88.2 – 91.2<br>(89.7 ± 1.3) | 0.7 – 1.5<br>(1.0 ± 0.4) | 3.3 – 6.7<br>(4.7 ± 1.2) |
| *D. pteropoda* | 5.0 – 7.8<br>(6.1 ± 0.4) | 2.4 – 5.2<br>(3.7 ± 1.0) | 344.3 – 373.7<br>(340.5 ± 12.3) | 390.2 – 400.7<br>(393.4 ± 4.1) | 26.0 – 54.3<br>(45.1 ± 3.5) | 90.4 – 92.8<br>(91.5 ± 1.0) | 0.9 – 2.0<br>(1.4 ± 0.4) | 2.3 – 4.2<br>(3.4 ± 0.7) |
| *D. sambiranensis*<br>subsp.<br>*bardotiae* | 2.6 – 6.1<br>(4.2 ± 0.2) | 3.5 – 8.6<br>(5.7 ± 1.0) | 353.8 – 366.3<br>(360.1 ± 3.4) | 380.2 – 393.2<br>(385.7 ± 4.6) | 39.9 – 58.3<br>(49.3 ± 2.2) | 84.2 – 91.6<br>(88.9 ± 2.9) | 0.7 – 1.0<br>(0.8 ± 0.1) | 2.6 – 6.2<br>(4.6 ± 1.2) |
| *D. sambiranensis*<br>subsp.<br>*sambiranensis* | 1.9 – 4.4<br>(3.1 ± 0.3) | 2.3 – 3.7<br>(3.0 ± 0.6) | 348.4 – 365.8<br>(356.7 ± 4.7) | 389.6 – 403.2<br>(393.1 ± 5.1) | 54.8 – 59.1<br>(57.8 ± 0.8) | 91.7 – 94.2<br>(92.9 ± 1.0) | 0.8 – 1.8<br>(1.0 ± 0.7) | 2.2 – 3.6<br>(3.0 ± 0.5) |
| *D. seriflora* | 4.0 – 7.3<br>(5.0 ± 0.3) | 2.2 – 5.0<br>(2.9 ± 1.0) | 352.3 – 376.2<br>(363.7 ± 6.5) | 386.1 – 398.1<br>(393.1 ± 4.1) | 51.6 – 61.8<br>(57.2 ± 1.5) | 90.1 – 93.9<br>(92.6 ± 1.4) | 0.7 – 1.5<br>(1.2 ± 0.4) | 2.3 – 4.3<br>(3.3 ± 0.7) |
| *Dioscorea*<br>species (Ovy<br>valiha) | 5.8 – 9.4<br>(6.9 ± 0.3) | 3.6 – 6.5<br>(4.6 ± 1.1) | 350.5 – 362.7<br>(357.0 ± 2.5) | 380.3 – 394.7<br>(386.1 ± 5.6) | 39.6 – 57.9<br>(51.8 ± 6.1) | 86.7 – 92.1<br>(89.7 ± 2.0) | 0.7 – 1.5<br>(1.0 ± 0.4) | 3.2 – 6.8<br>(4.7 ± 1.4) |

<sup>a</sup>Dried tubers consisting of parenchyma and periderm tissue. Protein content determined using the <sup>b</sup>Bradford and <sup>c</sup>Kjeldahl methods (n=10-16); energy content determined using the <sup>d</sup>ballistic bomb calorimeter and <sup>e</sup>Atwater methods (n=2-12); starch content determined using the <sup>f</sup>AOAC Official Method 996.11 for total starch determination (n=6-12); <sup>g</sup>total carbohydrate content determined by subtracting the protein, lipid, ash and water content (n=10-12); lipid content determined by <sup>h</sup>Folch method (n=10-12); and ash content determined by <sup>i</sup>mass after incineration (n=10-12). Data presented for the mean are shown as ± SEM.

### 3.2 Starch, carbohydrate, energy and lipid content

The nine wild edible *Dioscorea* species and cultivated *D. alata* from Madagascar were analysed over four harvest seasons for starch content, which was not consistently high or low for any one harvest season. The mean starch content of cultivated *D. alata* (57.8g/100g DW) across all four harvest seasons did not differ significantly from the mean starch content of the other edible species analysed (Table 2). The nine wild edible *Dioscorea* species and cultivated *D. alata* were analysed over six harvest seasons for total carbohydrate content, which ranged from 84.9g/100g (mean, DW) in cultivated *D. alata* to 92.9g/100g (mean, DW) in *D. sambiranensis* subsp. *sambiranensis* (Table 2). Total carbohydrate content in *D. alata* was significantly lower than that observed for *D.* sp. (Ovy valiha) (P<0.04), *D. pteropoda*, *D. sambiranensis* subsp. *sambiranensis*, *D. seriflora* (all P<0.01), *D. orangeana* (P<0.04) and *D. maciba* (P<0.04).

The nine wild edible yam species and cultivated *D. alata* from Madagascar were analysed for energy (kcal/100g) (Table 2). The mean energy provided by the edible yam species across all harvest seasons ranged from 340.4 kcal/100g (*D. buckleyana*) to 393.4 kcal/100g (*D. pteropoda*). The energy calculated for *D. alata* was significantly lower (P<0.01) than that observed for *D. pteropoda*, *D. sambiranensis* subsp. *sambiranensis* and *D. seriflora.* The total lipid content was calculated as 0.7 – 1.5g/100g (mean, DW) across all of the edible yam species analysed (Table 2), with the lipid content of all species analysed not significantly different to that observed for *D. alata*.

### 3.3 β-Carotene content

The edible yam species (wild and cultivated *D. alata*) were analysed for total carotenoid content over 8 harvest seasons between August 2018 and October 2019, with results expressed as beta-carotene equivalents (DW) (Figure 1). Differences in the β-carotene content of these species occurred between different harvest seasons, but the content was not consistently high or low for any one harvest season. This indicates that harvest season is not an informative indicator of β-carotene content in the edible yam species analysed.

**Figure 1.**
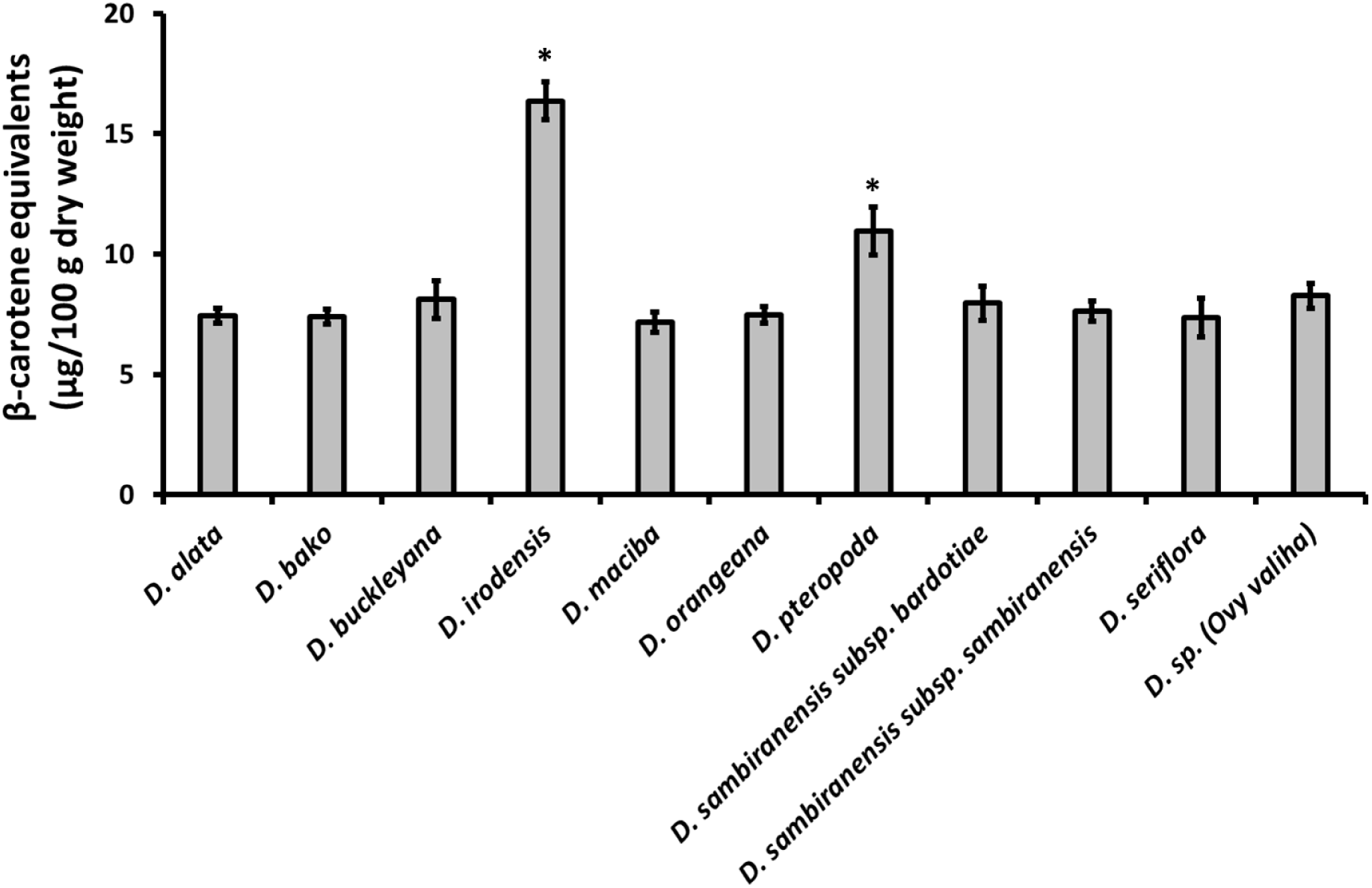
β-carotene content of nine wild edible yam species and cultivated *Dioscorea alata* across 8 harvest seasons, expressed as β-carotene equivalents (µg/100g dry weight) for mean values (n = 6 – 8) + SEM. *Significantly different from *D. alata* P<0.01.

The β-carotene content of *D. irodensis* was consistently higher than the β-carotene content of all other species analysed in all harvest seasons, except for the October 2019 harvest, when the β-carotene content of *D. pteropoda* (10.6µg/100g) and *D. sambiranensis* subsp*. bardotiae* (10.0µg/100g) was higher than that of *D. irodensis* (9.6µg/100g), but not significantly. The average (mean) β-carotene content of *D. irodensis* (16.4µg/100g) across all harvest seasons was significantly higher than the mean β-carotene content of all other species analysed, including *D. alata* (7.4µg/100g) (P<0.001; 1-way ANOVA and Tukey’s post-hoc test). The analysis also showed that the average β-carotene content of *D. pteropoda* was significantly higher than *D. alata* (P<0.02).

To assess the distribution of β-carotene in yam tuber parenchyma and periderm, the two species with the highest mean β-carotene content (*D. irodensis*:16.4µg/100g and *D. pteropoda*: 11.0µg/100g) were subjected to additional analysis for β-carotene content, compared with *D. alata*. The tuber parenchyma and periderm tissue of these three species across four harvest seasons (December 2019 – June 2020) was separated and each tissue was analysed separately to determine β-carotene content. β-Carotene was not significantly different between parenchyma and periderm tissue for each species analysed (Figure 2), indicating that peeling the tubers of these species to remove the periderm would not adversely affect the β-carotene content of the tubers (w/w). Consistent with the whole tuber β-carotene content (Figure 1), the mean β-carotene content of the separate parenchyma and periderm from *D. irodensis* (17.7 ± 0.6 and 18.0 ± 0.9 µg/100g, respectively; mean ± SEM, n = 12) was significantly higher than the β-carotene content of *D. pteropoda* (14.4 ± 0.8 and 13.2 ± 0.5 µg/100g, respectively; mean ± SEM, n = 12) and *D. alata* (12.2 ± 0.4 and 11.9 ± 0.5 µg/100g, respectively; mean ± SEM, n = 12) (parenchyma, P<0.001; periderm, P<0.001; 1-way ANOVA and Tukey’s post-hoc test). The β-carotene content of *D. irodensis* was also significantly higher than *D. alata* for both parenchyma and periderm in each harvest season (Figure 2).

**Figure 2.**
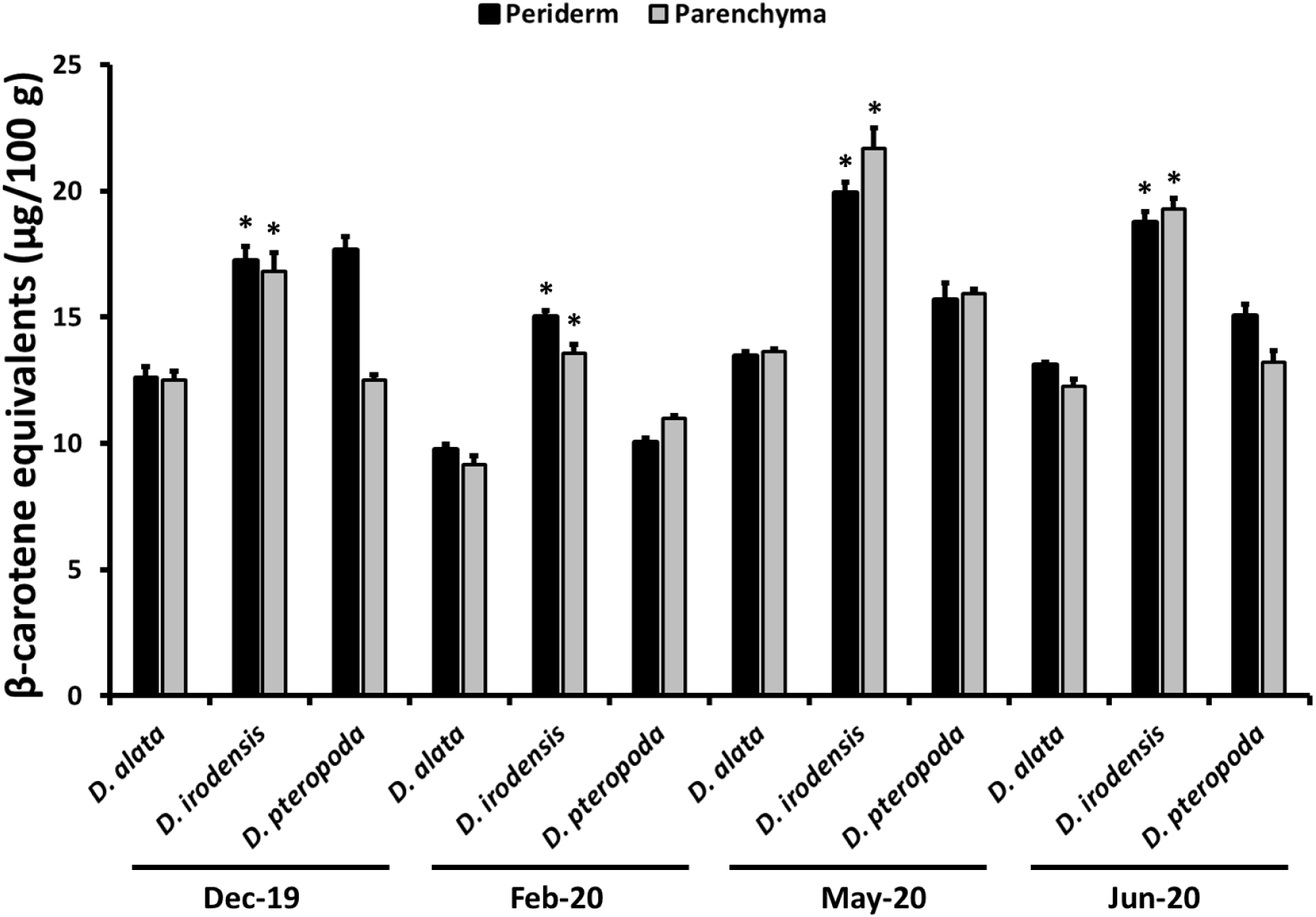
β-carotene content of separated parenchyma and periderm tissue of two wild edible yam species and cultivated *Dioscorea alata* across 4 harvest seasons, expressed as β-carotene equivalents (µg/100g dry weight) for mean values (n = 3) + SEM. *Significantly different from *D. alata* in each harvest season, P<0.01.

### 3.4 Mineral and ash content

The total ash content, representing the inorganic, or crude mineral, content determined for the edible yam species analysed in this study was in the range of 3.0 – 6.0g/100g (mean; DW). The total ash content of *D. alata* was significantly higher than that observed in *D. pteropoda (*P<0.03), *D. sambiranensis* subsp. *sambiranensis* (P<0.01) and *D. seriflora* (P<0.02). The wild edible yam species and cultivated *D. alata* were also analysed for calcium, iron and zinc content over four harvest seasons between August 2018 and March 2019, with (DW) content determined by ICP-MS. Differences in the content of these minerals occurred between different harvest seasons, but the content was not consistently high or low for any one harvest season. This indicates that harvest season is not an informative indicator of calcium, iron or zinc content in the edible yam species analysed.

Although there were no statistical differences in mean mineral content across these four harvest seasons, the mean calcium content of *D. buckleyana* (2456mg/kg), *D. irodensis* (2354mg/kg), and *D. sambiranensis* subsp. *bardotiae* (2346mg/kg) was 2.8 – 3 times greater than *D. alata* (833mg/kg) (Figure 3A). For iron, the content in *D. alata* (58mg/kg) was lower than the mean iron content of all other species analysed; the highest levels were found in *D. irodensis* (311mg/kg) (Figure 3B). The mean zinc content of *D. alata* (35mg/kg) across the same period was higher than the mean zinc content of all other species analysed, although not significantly (Figure 3C).

**Figure 3.**
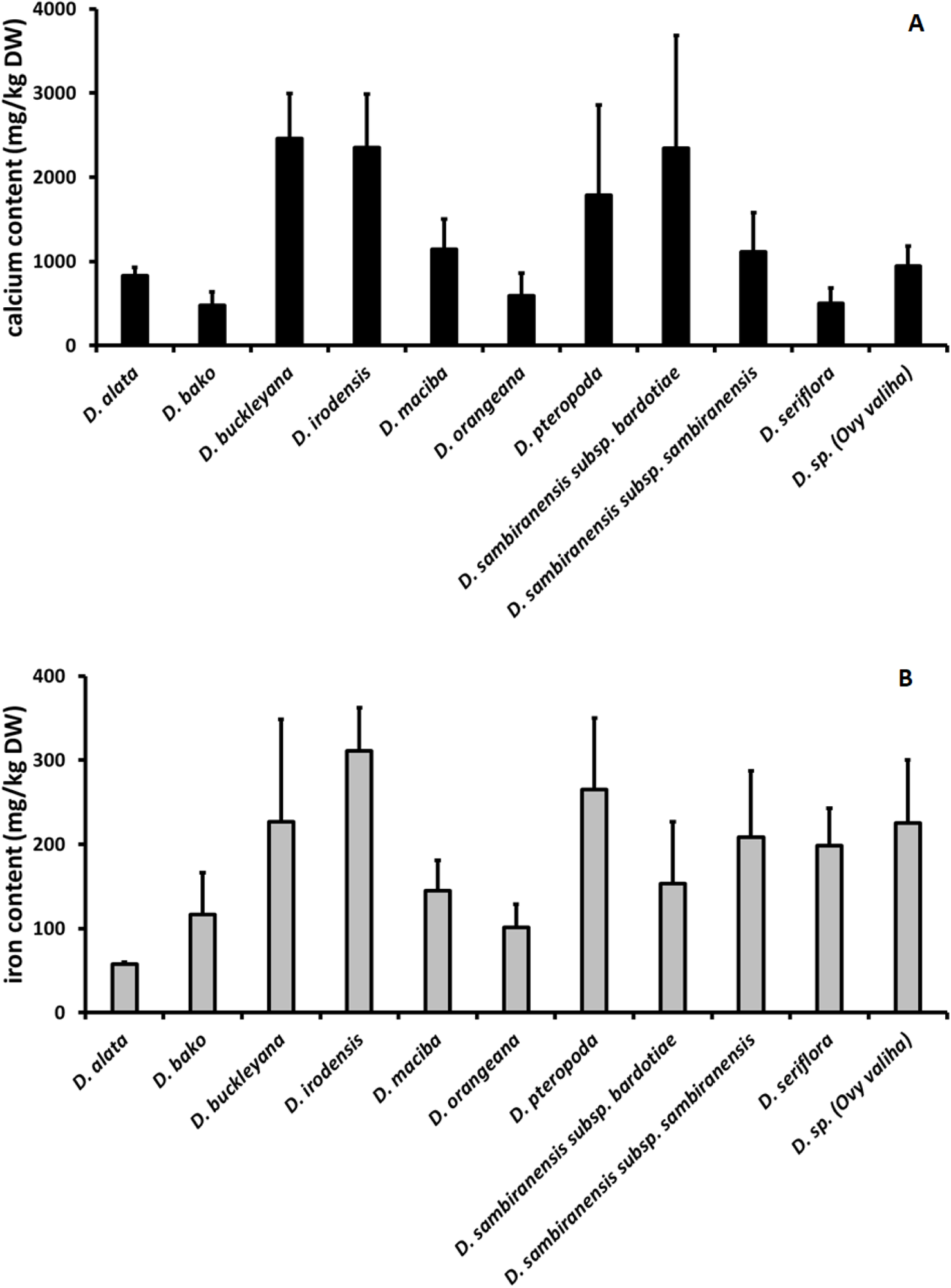
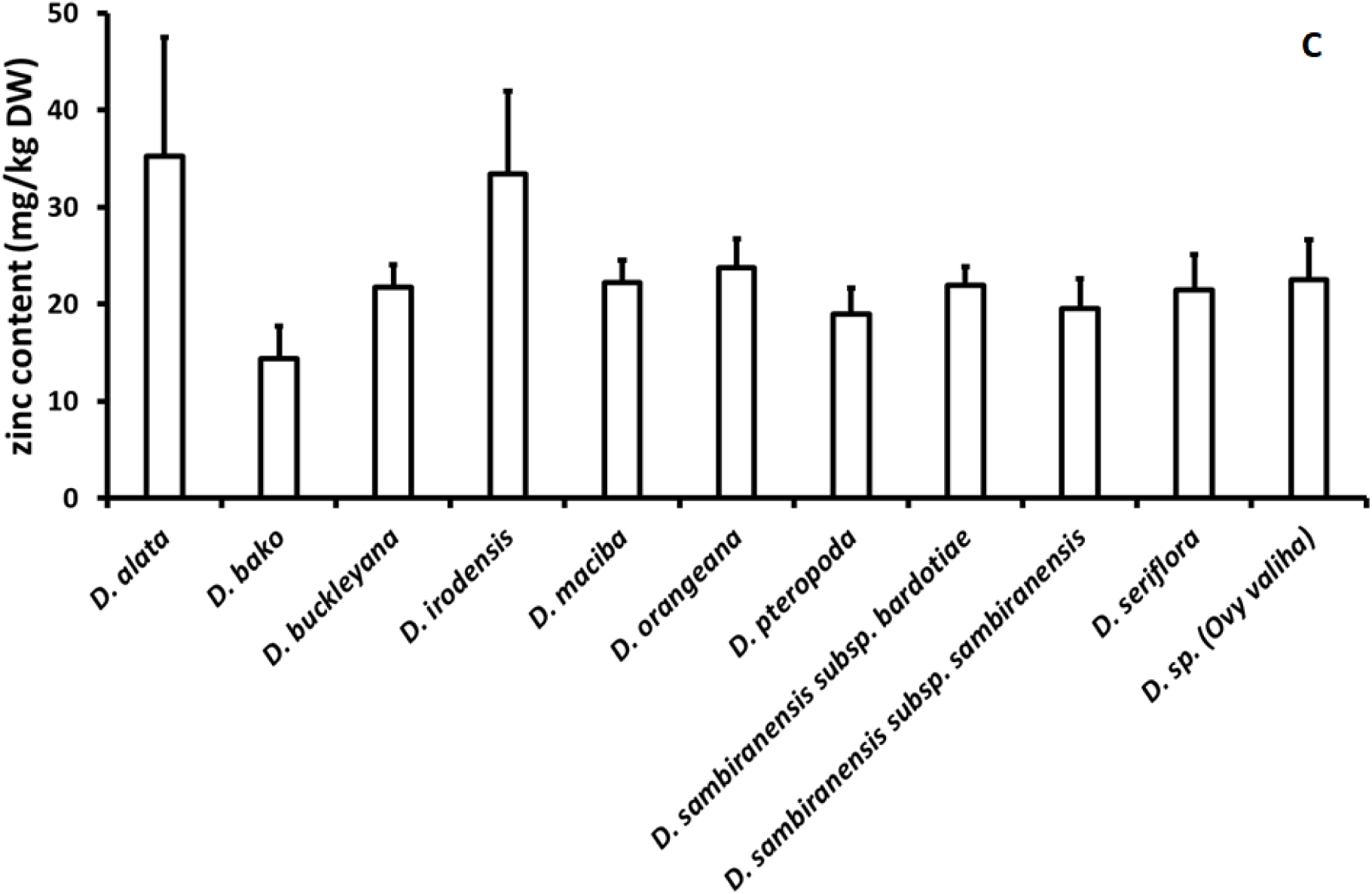
Mineral (A: calcium, B: iron, C: zinc) content of nine wild edible yam species and cultivated *Dioscorea alata* across 4 harvest seasons. (n = 3 – 4) + SEM.

## Discussion

In Madagascar, wild edible yams are consumed in addition to the non-endemic cultivated winged yam, *Dioscorea alata*. However, the nutrient composition of wild edible yams (an important Malagasy food resource, particularly in times of hardship and famine) has been understudied. Consequently, there is a need to further understand how wild edible yams may contribute to dietary nutrition in Madagascar and how their nutrient profiles compare with the cultivated *D. alata*. This study addresses this knowledge gap by presenting new nutrient composition data for nine wild edible *Dioscorea* species in comparison with cultivated *D. alata*, also sourced from Madagascar.

Dietary protein is required for optimal growth, development and health in humans. Although protein from animal sources is considered to provide higher quantities and more balanced proportions of amino acids compared to protein from plant sources, plant-based foods contribute 35% of protein globally to human diets, while consumption of <65% of total protein from animal-sourced foods can lead to protein under-nutrition (Wu, 2016). In this context, plant-based protein can contribute to dietary protein requirements, particularly in circumstances or periods where access to other food sources of protein may be limited. Indeed, although yams have been primarily considered to be a starchy crop, they may also contribute to dietary protein, as has been documented for edible yams in West Africa (Asiedu and Sartie, 2010). The protein content of cultivated *D. alata* was analysed and compared with the protein content of the nine edible wild yam species in this study. The mean protein content of cultivated *D. alata* did not differ significantly from the mean protein content of any of the wild edible species analysed (tubers harvested 2018-2019). In tubers harvested in 2019-2020, the protein content of *D. alata* was significantly higher than all wild edible species analysed, except for *D. irodensis*, *D. sambiranensis* subsp. *bardotiae* and *D. bako.* These results indicate that none of the wild edible yams analysed in this study are a better source of protein in comparison with cultivated *D. alata*, but certain species may provide comparable protein levels to those provided by *D. alata*. However, it is also recognised that the amino acid profiles of the protein occurring in *D. alata* and the wild edible yams require analysis to further evaluate their contribution as plant-based protein sources for dietary needs.

The observed protein content of *D. alata* and all other edible wild yam species analysed in this study (range for mean values: 2.9 – 8.5% DW) compares favourably with 2.4% – 7.8% protein (DW) reported for different cultivars of sweet potato (Kourouma et al., 2020) [although other studies report a lower range: 0.6 – 2.5% (DW) (Neela and Fanta, 2019)], and is higher than protein reported for cassava (1 – 3% DW) (Ferraro et al., 2016; Shewry, 2003). Cultivated *D. alata* and wild edible yams in Madagascar may therefore be a richer protein source protein compared to some other tuber crops such as cassava.

Starch forms the bulk of energy provided by edible *Dioscorea* species. For example, *D. esculenta* (Lour.) Burkill tubers have been shown to contain similar levels of starch (78% DW) compared to *D. alata* (65 - 84% starch, tubers not sourced from Madagascar) (Wilkin et al., 2022). In this study, the mean starch content of cultivated *D. alata* across four harvest seasons did not differ significantly from the mean starch content of the wild edible *Dioscorea* species analysed. The edible *Dioscorea* species were also analysed over six harvest seasons for total carbohydrate content, which ranged from 84.9g/100g (mean, DW) in cultivated *D. alata* to 92.9g/100g (mean, DW) in *D. sambiranensis* subsp. *sambiranensis*. Total carbohydrate in *D. alata* was significantly lower than in *D.* sp. (Ovy valiha), *D. pteropoda*, *D. sambiranensis* subsp. *sambiranensis*, *D. seriflora*, *D. orangeana* and *D. maciba.* These results indicate that *D. alata* has comparable starch levels to the wild edible *Dioscorea* species analysed, while certain wild edible species may provide a higher total carbohydrate content (which represents starch and individual sugars) compared to *D. alata*. These results compare with 80 – 90g/100g (DW) carbohydrate, of which 80g/100g is starch, for cassava root (Ferraro et al., 2016); and 51.9 – 69.2g/100g (DW) starch content which has been reported for different cultivars of sweet potato (Kourouma et al., 2020), while the total carbohydrate for different varieties of sweet potato is reported as 81.3 – 87.3g/100g (DW) (Cartier et al., 2017). Thus, *D. alata* and the wild edible yams analysed in this study have a comparable total carbohydrate and starch content to the tuber crops sweet potato and cassava.

Yams are a major source of calories for millions of people in tropical and sub-tropical regions (Asiedu and Sartie, 2010). In this study, the calculated energy for *D. alata* was significantly lower than that calculated for *D. pteropoda*, *D. sambiranensis* subsp. *sambiranensis* and *D. seriflora*. However, the energy levels observed for *D. alata* and the wild edible *Dioscorea* species analysed are comparable to energy values for orange-fleshed sweet potato (345 kcal/100g DW) (Neela and Fanta, 2019). Furthermore, the total lipid content across all of the edible yam species analysed (which did not differ significantly between species) was comparable with orange-fleshed sweet potato (0.03 – 0.9 g/100g DW) (Neela and Fanta, 2019) and with other cultivars of sweet potato (0.24 – 1.11g/100g DW) (Kourouma et al., 2020). However, there are some reports of lipid content of sweet potato being as high as 2.2g/100g (Cartier et al., 2017).

Vitamin A is a fat-soluble vitamin that occurs in the diet in various forms (e.g. as retinol isomers in animal products), depending on the source. It is essential for growth, development and maintenance of epithelial tissue and for vision, while a deficiency can result in ocular complications (e.g. xeropthalmia which may progress to blindness) (Martindale, 2022). Certain edible plants, including various fruits and vegetables, are a source of provitamin A carotenoids, which have vitamin A activity when consumed, with β-carotene being the most important (McCance and Widdowson, 2015). To understand the contribution of the nine wild edible yams and the cultivated *D. alata* from Madagascar have on dietary vitamin A intake, they were assessed for their carotenoid content over eight harvest seasons, with results expressed as β-carotene equivalents (DW). The β-carotene content of *D. irodensis* was consistently higher than the β-carotene content of all other species analysed in all harvest seasons and the average (mean) β-carotene content of *D. irodensis* across all harvest seasons was significantly higher than the mean β-carotene content of all other species analysed, including the cultivated *D. alata*.

To assess the distribution of β-carotene in yam tuber parenchyma (which is consumed by people) and periderm (which is not consumed by people but may be used as livestock feed), the two species with the highest mean β-carotene content (*D. irodensis* and *D. pteropoda*) and *D. alata* were subjected to additional analysis for β-carotene content in the parenchyma and periderm tissue. β-Carotene content was not significantly different between parenchyma and periderm tissue for each species analysed (Figure 2), indicating that peeling the tubers of these species to remove the periderm would not adversely affect the β-carotene content of the tubers (w/w), and that the inclusion of periderm tissue in livestock feed could provide a source of β-carotene (albeit at low levels). Consistent with the whole tuber β-carotene content (Figure 1), the mean β-carotene content of the separate parenchyma and periderm from *D. irodensis* was also significantly higher than that in *D. alata*.

Food tables indicate that yam contains only trace amounts of carotenoids (McCance and Widdowson, 2015). Our data showing β-carotene equivalents in the low µg range are consistent with food table values and contrast with 416µg/100g – 20,811µg/100g (DW) β-carotene reported for different cultivars of sweet potato (Kourouma et al., 2020). Although the β-carotene content of *D. irodensis* was higher than all other edible yam species analysed in this study, this value is still at least 25-fold less than that observed for certain sweet potato varieties (Kourouma et al., 2020). Therefore, considering the low β-carotene content compared to certain other edible plants such as sweet potatoes, consumption of cultivated *D. alata* or the wild edible yam species investigated in this study, are unlikely to provide a meaningful contribution to dietary vitamin A intake or to prevent vitamin A deficiency. It should also be considered, however, that carotenoids may have functions beyond their role in nutrition as vitamin A precursors. Dietary intake of carotenoids has been associated with a reduced risk of certain chronic diseases and increased mechanistic effects (e.g. antioxidant) relevant to human health (Monjotin et al., 2022; Bohn et al., 2021). Therefore, although the β-carotene content of the wild and cultivated yams from Madagascar are unlikely to significantly contribute to dietary vitamin A requirements, the other functions of carotenoids, and those of other phytochemical constituents, merit further consideration for other potential health benefits.

The total ash content, representing the inorganic, or crude mineral, content determined for the edible yam species analysed in this study ranged from 3.0 – 6.0g/100g (mean; DW), which compares similarly with that observed for different sweet potato cultivars (2.0 – 4.5g/100g DW) (Kourouma et al., 2020; Cartier et al., 2017). The wild edible yam species and cultivated *D. alata* were also analysed for calcium, iron and zinc content over four harvest seasons. Although there were no statistical differences in mean mineral content across these four harvest seasons, the mean calcium content of *D. buckleyana*, *D. irodensis*, and *D. sambiranensis* subsp. *bardotiae* was 2.8 – 3 times greater than *D. alata* (Figure 3A). For iron, the content in *D. irodensis* was 5 times higher than in *D. alata* (Figure 3B). Analysis showed that mean zinc content was similar across all yam species (Figure 3C).

All mineral values were substantially higher than those reported for raw yam tubers in food tables (McCance and Widdowson, 2015), which may indicate that a significant proportion of the mineral content lies in the periderm. The difference in mineral content observed for different collections of the same species could be explained by different environmental conditions i.e. soil fertility and climate of the region, which may be associated with the origin or time of harvest (Huang et al., 2007). Many studies on wild edible plant species show that they contain a considerable percentage of minerals, ranging from 2.48 to 6.36% (Afiukwa et al., 2013; Oko & Famurewa, 2015; Shajeela et al., 2011), which is comparable to the observations in our study. These findings, therefore, further support other research that shows reduced access to wild foods can negatively affect food security and nutrition, specifically micronutrient consumption (Moore et al., 2022).

Different varieties of orange-fleshed sweet potato contain calcium in the range 219.8 to 273.5mg/kg (DW), iron in the range 9.1 to 14.0mg/kg (DW) and zinc in the range 28.5 to 42.4mg/kg (DW) (Alam et al., 2020), while other varieties improved for disease tolerance are reported to contain calcium, iron and zinc in the ranges 207.0 – 250.9, 50.9 – 101.8 and 21.8 – 31.8mg/kg (DW), respectively (Mitiku and Teka, 2017). The mineral content of the tubers of different cassava (yellow-fleshed) genotypes is reported as 1149.7 – 1444.7mg/kg (DW) for calcium, 8.5 – 9.5mg/kg (DW) for iron, and 8.7 – 9.7mg/kg (DW) for zinc (Alamu et al., 2020). Other cassava genotypes are reported to contain 1360 – 3690mg/kg (DW) calcium, 290 – 400mg/kg (DW) iron and 130 – 190mg/kg (DW) zinc (Charles et al., 2005). Thus, the calcium content of wild yams and cultivated *D. alata* observed in this study are higher than calcium levels reported for certain varieties of sweet potato, and within a similar range to those reported for certain genotypes of cassava. The iron and zinc contents observed for wild yams and cultivated *D. alata* were within the content range of these minerals that are reported to occur in sweet potato and cassava tubers, depending on the variety or genotype of these tuber crops. An important caveat is that the reported mineral values represent the content of the raw plant tissue. To date there have been no reports of the bioavailability of these minerals (i.e. the fraction that can be absorbed in the intestine and utilised in the body) from either raw material or from meals containing cooked yam.

Knowledge of the nutrient composition of wild edible yams in Madagascar may have different implications for food security and conservation of edible species. Firstly, the knowledge that certain wild edible yams contain comparable nutrient contents (e.g. protein, starch) to the cultivated *D. alata* could potentially enhance the demand for wild edible yams, particularly when cultivated crops (*D. alata* and others) may be in limited supply. Secondly, although wild edible yams are harvested in certain regions of Madagascar to supplement crops such as cassava and maize, especially during hunger periods (Andriamparany et al., 2014), the knowledge that wild edible yams may contain a higher protein content than other crops such as cassava (1 – 3% (DW) (Ferraro et al., 2016; Shewry, 2003)) may provide an additional incentive to harvest wild edible yams during other periods, exacerbating existing threats to their survival. Overharvesting of edible plants from the wild may have consequences for their conservation and for food security. Indeed, certain *Dioscorea* species are threatened and overharvesting of tubers from the wild is a contributing factor, with some species already categorised as Endangered according to the IUCN Red List (Table 1). However, although wild edible species may be exploited for their useful properties, knowledge of how they may benefit people, for example, as food or medicines, may also motivate the sustainable management of natural resources (Howes et al., 2020). From this perspective, knowledge of the nutrient content of wild edible food species could provide incentives for their sustainable cultivation and use, to align with strategies for their conservation and for food security, in addition to supporting livelihoods. It is recognised, however, that other factors are to be considered when selecting edible yams to harvest from the wild, or indeed to cultivate. For example, local use of wild edible yams may depend on taste or personal preferences, local needs, availability, market prices, location and quantities harvested (Andriamparany et al., 2014) amongst other complexities, as has been discussed previously (Moore et al., 2022). In this study, the wild edible yams analysed include 6/6 (*D. bako*, *D. buckleyana*, *D. maciba*, *D. orangeana*, *D. sambiranensis*, *D. seriflora*) of the most favoured wild edible yam species in Madagascar (Wilkin et al., 2022). Here, we therefore present new nutrient composition data for the six most favoured wild edible yams in Madagascar, to complement other food research studies on factors that influence food choice (e.g. taste, texture), while providing new nutrient data for popular edible yams in Madagascar, relevant to help support their sustainable use and conservation.

As a small island, Madagascar is vulnerable to the impacts of climate change on food and nutrition security due to factors such as fragile natural environments and lack of arable land, impacting on agriculture and contributing to risks including water insecurity (WHO and UN, 2021). Furthermore, climate change is likely to exacerbate the triple burden of metabolic and lifestyle risk factors for non-communicable diseases and malnutrition (WHO and UN, 2021), in a country that already has the highest rate of chronic malnutrition in the world, with 42% of children under age five suffering from stunting (Moore et al., 2022). Potential strategies to ameliorate the effects of climate change on food security and dietary diversity include creating more diverse and climate-resilient agricultural production systems in addition to improved knowledge and cultivation of naturally stress-resistant plants, aligned with methods to maintain the genetic diversity of crops (Ulian et al., 2020; Herrera et al., 2021). Another approach to address these challenges is to promote the cultivation of nutrient-dense indigenous vegetables, rather than more recently introduced vegetables that may not be adapted to local growing conditions or be culturally suitable; for example, nonprofit organizations in Madagascar have aided the cultivation of indigenous yams (Moore et al., 2022). Furthermore, community involvement in conservation projects has greater potential to achieve positive outcomes for both biodiversity conservation and food security (Moore et al., 2022). In this context, dissemination of information on the nutrient content of wild edible plants, such as for wild edible yams in Madagascar as presented in this study, could facilitate motivation and opportunities for the sustainable cultivation and use of endemic yams to benefit people and the environment. Finally, it is recognized that although new nutrient composition data for wild edible yams from Madagascar is a first step to understanding how these food resources may contribute to dietary nutrient requirements, this knowledge can also underpin future studies to further evaluate the role of wild edible yams in the diet, including research to assess the effects of preparation and cooking on nutrient content, and the bioavailability and bioaccessibility of nutrients.

## Conclusion

This study presents new information on the nutrient composition of nine edible yams (*Dioscorea* species) endemic to Madagascar, presenting new data for their protein, carbohydrate/starch, mineral and lipid content, as well as the energy they provide. The results show that although none of the wild edible yams analysed in this study are a better source of protein compared to cultivated *D. alata*, all of the *Dioscorea* species analysed contain higher levels of protein compared to those published for cassava. The wild edible *D. irodensis* contains a significantly higher β-carotene content compared to all other edible yams analysed, but levels are still considerably lower (at least 25-fold) than published data for certain sweet potato varieties. This study has also revealed that of the wild edible *Dioscorea* species analysed, *D. buckleyana*, *D. irodensis* and *D. sambiranensis* subsp. *bardotiae* have a higher calcium content compared to the cultivated *D. alata*, while all nine wild edible yam species contain higher iron content, compared to cultivated *D. alata*. In conclusion, these results provide new insights into the nutrient composition of wild edible yams endemic to Madagascar which could underpin the direction of future research to further understanding of how they may contribute to dietary nutritional requirements and dietary diversity in Madagascar. Dissemination of new knowledge on the nutritional value of wild edible yams could also facilitate opportunities to incentivise their conservation, sustainable cultivation and use, to complement strategies to enhance food security in Madagascar.

## Acknowledgements

The authors thank the Darwin Initiative for funding the following projects to support this research: ‘Sustainable yam markets for conservation and food security in Madagascar’ (EIDPO049) and ‘Conserving Madagascar’s yams through cultivation for livelihoods and food security’ (22-005).

