## Supplementary Table 1 for "Wild edible yams from Madagascar: New insights into nutritional composition support their use for food security and conservation"

**Supplementary Table 1.** Collection details for edible yam tubers in Madagascar.

| ***Dioscorea* species** | **Harvest month and year** | **Harvest location (GPS)** | **Collection reference codes** |
|---|---|---|---|
| *D. bako* | August 2018 | Lat 19.9582/ Long 44.6487 | MOR 01<br>BI 28865 |
| *D. buckleyana* | August 2018 | Lat 12.2125/ Long 49.200 | FEN 409 (M22)<br>BI 28866 |
| *D. maciba* | August 2018 | Lat 12.2449/ Long 49.2144 | FEN 407 (M42)<br>BI 28867 |
| *D. orangeana* | August 2018 | Lat 12.1938/ Long 49.2435 | FEN 411 (M52)<br>BI 28868 |
| *D. pteropoda* | August 2018 | Lat 12.2449/ Long 49.2508 | FEN 406 (M21)<br>BI 28869 |
| *D. sambiranensis* subsp. *bardotiae* | August 2018 | Lat 12.2117/ Long 49.2023 | FEN 408 (M12)<br>BI 28870 |
| *D. sambiranensis* subsp. *sambiranensis* | August 2018 | Lat 12.2436/ Long 49.2018 | FEN 413 (M30)<br>BI 28871 |
| *D. seriflora* | August 2018 | Lat 13.4744/ Long 48.3004 | FEN 445 (M51)<br>BI 28872 |
| *Dioscorea* species (Ovy valiha) | August 2018 | Lat 12.3729/ Long 49.2447 | FEN 415 (M11)<br>BI 28873 |
| *D. alata* | October 2018 | Lat 13.4740/ Long 48.2943 | FEN 479 (M33)<br>BI 29015 |
| *D. bako* | October 2018 | Lat 19.7762/ Long 44.8202 | MOR 02 (M61)<br>BI 29012 |
| *D. buckleyana* | October 2018 | Lat 12.2125/ Long 49.2001 | FEN 473 (M27)<br>BI 29107 |
| *D. irodensis* | October 2018 | Lat 12.1836/ Long 49.2245 | FEN 475 (M46)<br>BI 29008 |
| *D. maciba* | October 2018 | Lat 12.2429/ Long 49.2144 | FEN 471 (M47)<br>BI 29007 |
| *D. orangeana* | October 2018 | Lat 12.1938/ Long 49.2435 | FEN 474 (M57)<br>BI 29011 |
| *D. pteropoda* | October 2018 | Lat 12.2449/ Long 49.2508 | FEN 476 (M26)<br>BI 29013 |
| *D. sambiranensis* subsp. *bardotiae* | October 2018 | Lat 12.2117/ Long 49.2023 | FEN 472 (M17)<br>BI 29010 |
| *D. sambiranensis* subsp. *sambiranensis* | October 2018 | Lat 13.2842/ Long 48.4528 | FEN 480 (M35)<br>BI 29009 |
| *D. seriflora* | October 2018 | Lat 12.4744/ Long 49.3004 | FEN 478 (M56) |

| | | | |
|---|---|---|---|
| | | | BI 29016 |
| *Dioscorea* species (Ovy valiha) | October 2018 | Lat 12.3729/ Long 49.2447 | FEN 477 (M16)<br>BI 29014 |
| *D. alata* | December 2018 | Lat 13.4740/ Long 48.2943 | FEN 521 (M38)<br>BI 29122 |
| *D. bako* | December 2018 | Lat 19.7762/ Long 44.8202 | MOR 03<br>BI 29123 |
| *D. buckleyana* | December 2018 | Lat 12.2125/ Long 49.2001 | FEN 505 (M24)<br>BI 28124 |
| *D. irodensis* | December 2018 | Lat 12.1836/ Long 49.2245 | FEN 507 (M43)<br>BI 29126 |
| *D. maciba* | December 2018 | Lat 12.2429/ Long 49.2144 | FEN 508 (M44)<br>BI 29127 |
| *D. orangeana* | December 2018 | Lat 12.1938/ Long 49.2435 | FEN 506 (M54)<br>BI 299128 |
| *D. pteropoda* | December 2018 | Lat 12.2449/ Long 49.2508 | FEN 509 (M23)<br>BI 29125 |
| *D. sambiranensis* subsp. *bardotiae* | December 2018 | Lat 12.2117/ Long 49.2023 | FEN 504 (M14)<br>BI 29129 |
| *D. sambiranensis* subsp. *sambiranensis* | December 2018 | Lat 13.2842/ Long 48.4528 | FEN 522<br>BI 29130 |
| *D. seriflora* | December 2018 | Lat 12.4744/ Long 49.3004 | FEN 520<br>BI 29131 |
| *Dioscorea* species (Ovy valiha) | December 2018 | Lat 12.3729/ Long 49.2447 | FEN 510<br>BI 29132 |
| *D. bako* | March 2019 | Lat 19.7762/ Long 44.8202 | MOR 04<br>BI 29272 |
| *D. buckleyana* | March 2019 | Lat 12.2125/ Long 49.2001 | GHM 78<br>BI 29272 |
| *D. irodensis* | March 2019 | Lat 12.1836/ Long 49.2245 | GHM 80<br>BI 29274 |
| *D. maciba* | March 2019 | Lat 12.2429/ Long 49.2144 | GHM 82<br>BI 29275 |
| *D. orangeana* | March 2019 | Lat 12.1938/ Long 49.2435 | GHM 79<br>BI 29276 |
| *D. pteropoda* | March 2019 | Lat 12.2449/ Long 49.2508 | GHM 81<br>BI 29277 |
| *D. sambiranensis* subsp. *bardotiae* | March 2019 | Lat 12.2117/ Long 49.2023 | GHM 77<br>BI 29278 |
| *D. sambiranensis* subsp. *sambiranensis* | March 2019 | Lat 13.2842/ Long 48.4528 | GHM 74 |

| | | | |
|---|---|---|---|
| | | | BI 29279 |
| *D. seriflora* | March 2019 | Lat 12.4744/ Long 49.3004 | GHM 75<br>BI 29280 |
| *Dioscorea* species (Ovy valiha) | March 2019 | Lat 12.3729/ Long 49.2447 | GHM 76<br>BI 29281 |
| *D. alata* | April 2019 | Lat 13.4740/ Long 48.2943 | GHM 122<br>BI 29753 |
| *D. bako* | April 2019 | Lat 19.9562/ Long 44.65033 | MOR 05<br>BI 29754 |
| *D. buckleyana* | April 2019 | Lat 12.2125/ Long 49.2001 | GHM 126<br>BI 29755 |
| *D. irodensis* | April 2019 | Lat 12.1836/ Long 49.2245 | FEN 709<br>BI 29756 |
| *D. maciba* | April 2019 | Lat 12.2429/ Long 49.2144 | GHM 124<br>BI 29757 |
| *D. orangeana* | April 2019 | Lat 12.1938/ Long 49.2435 | FEN 708<br>BI 29758 |
| *D. pteropoda* | April 2019 | Lat 12.2449/ Long 49.2508 | FEN 707<br>BI 29755 |
| *D. sambiranensis* subsp. *bardotiae* | April 2019 | Lat 12.2117/ Long 49.2023 | GHM 125<br>BI 29760 |
| *D. sambiranensis* subsp. *sambiranensis* | April 2019 | Lat 13.2842/ Long 48.4528 | FEN 706<br>BI 29761 |
| *D. seriflora* | April 2019 | Lat 12.4744/ Long 49.3004 | GHM 123<br>BI 29762 |
| *Dioscorea* species (Ovy valiha) | April 2019 | Lat 12.3729/ Long 49.2447 | FEN 710<br>BI 29763 |
| *D. alata* | June 2019 | Lat 13.4740/ Long 48.2943 | MIR125<br>BI 30899 |
| *D. bako* | June 2019 | Lat 19.7762/ Long 44.8202 | MOR 06<br>BI 30902 |
| *D. buckleyana* | June 2019 | Lat 12.2125/ Long 49.2003 | GHM184<br>BI 30895 |
| *D. irodensis* | June 2019 | Lat 12.1837/ Long 49.2244 | GHM182<br>BI 30901 |
| *D. maciba* | June 2019 | Lat 12.2430/ Long 49.2145 | GHM180<br>BI 30896 |
| *D. orangeana* | June 2019 | Lat 12.1938/ Long 49.2437 | GHM181<br>BI 30900 |
| *D. pteropoda* | June 2019 | Lat 12.2448/ Long 49.2509 | GHM179 |

| | | | |
|---|---|---|---|
| | | | BI 30894 |
| *D. sambiranensis* subsp. *bardotiae* | June 2019 | Lat 12.2118/ Long 49.2024 | GHM183<br>BI 30897 |
| *D. sambiranensis* subsp. *sambiranensis* | June 2019 | Lat 13.2842/ Long 48.4530 | MIR126<br>BI 30898 |
| *D. seriflora* | June 2019 | Lat 12.4745/ Long 48.3004 | MIR124<br>BI 30903 |
| *Dioscorea* species (Ovy valiha) | June 2019 | Lat 12.3728 Long 49.2448 | GHM178<br>BI 30893 |
| *D. alata* | August 2019 | Lat 13.4740/ Long 48.2943 | GHM188<br>BI 30910 |
| *D. bako* | August 2019 | Lat 19.7642/ Long 44.6516 | MOR 07<br>BI 30913 |
| *D. buckleyana* | August 2019 | Lat 12.2005/ Long 49.2228 | MIR151<br>BI 30906 |
| *D. irodensis* | August 2019 | Lat 12.3422/ Long 49.2436 | GHM185<br>BI 30912 |
| *D. maciba* | August 2019 | Lat 12.2346/ Long 49.2089 | GHM187<br>BI 30907 |
| *D. orangeana* | August 2019 | Lat 12.1917/ Long 49.2002 | FEN805<br>BI 30911 |
| *D. pteropoda* | August 2019 | Lat 12.2448/ Long 49.2508 | GHM186<br>BI 30905 |
| *D. sambiranensis* subsp. *bardotiae* | August 2019 | Lat 12.2233/ Long 49.1929 | FEN806<br>BI 30908 |
| *D. sambiranensis* subsp. *sambiranensis* | August 2019 | Lat 13.2843/ Long 48.4526 | MIR152<br>BI 30909 |
| *D. seriflora* | August 2019 | Lat 13.4745/ Long 48.0841 | FEN807<br>BI 30914 |
| *Dioscorea* species (Ovy valiha) | August 2019 | Lat 12.3729/ Long 49.2449 | FEN804<br>BI 30904 |
| *D. alata* | October 2019 | Lat 13.4740/ Long 48.2943 | FEN809<br>BI 30921 |
| *D. bako* | October 2019 | Lat 19.7642/ Long 44.6516 | MOR 10<br>BI 30924 |
| *D. buckleyana* | October 2019 | Lat 12.2005/ Long 49.2228 | GHM222<br>BI 30917 |
| *D. irodensis* | October 2019 | Lat 12.3422/ Long 49.2436 | GHM224<br>BI 30923 |
| *D. maciba* | October 2019 | Lat 12.2346/ Long 49.2089 | GHM223 |

| | | | |
|---|---|---|---|
| | | | BI 30918 |
| *D. orangeana* | October 2019 | Lat 12.1917/ Long 49.2002 | GHM221<br>BI 30922 |
| *D. pteropoda* | October 2019 | Lat 12.2448/ Long 49.2508 | MIR170<br>BI 30916 |
| *D. sambiranensis* subsp. *bardotiae* | October 2019 | Lat 12.2233/ Long 49.1929 | FEN810<br>BI 30919 |
| *D. sambiranensis* subsp. *sambiranensis* | October 2019 | Lat 13.2843/ Long 48.4526 | GHM220<br>BI 30920 |
| *D. seriflora* | October 2019 | Lat 13.4948/ Long 48.0841 | FEN808<br>BI 30925 |
| *Dioscorea* species (Ovy valiha) | October 2019 | Lat 12.3729/ Long 49.2449 | FEN811<br>BI 30915 |
| *D. alata* | December 2019 | Lat 13.4740/ Long 48.2943 | FEN826<br>BI 31139[a] / BI 31140[b] |
| *D. bako* | December 2019 | Lat 19.77318611/ Long 44.83166667 | MOR11<br>BI 31119[a] / BI 31120[b] |
| *D. buckleyana* | December 2019 | Lat 12.2004/ Long 49.2204 | MIR174<br>BI 31123[a] / BI 311124[b] |
| *D. irodensis* | December 2019 | Lat 12.3419/ Long 49.2447 | FEN829<br>BI 31125[a] / BI 31126[b] |
| *D. maciba* | December 2019 | Lat 12.2330/ Long 49.1945 | MIR172<br>BI 31129[a] / BI 31130[b] |
| *D. orangeana* | December 2019 | Lat 12.1917/<br>Long 49.2001 | MIR175<br>BI 31121[a] / BI 31122[b] |
| *D. pteropoda* | December 2019 | Lat 12.2450/ Long 49.2515 | MIR176<br>BI 31137[a] / BI 31138[b] |
| *D. sambiranensis* subsp. *bardotiae* | December 2019 | Lat 12.2222/ Long 49.1936 | MIR173<br>BI 31127[a] / BI 31128[b] |
| *D. sambiranensis* subsp. *sambiranensis* | December 2019 | Lat 13.2845/ Long 48.4523 | FENj827<br>BI 31133[a] / BI 31134[b] |
| *D. seriflora* | December 2019 | Lat 13.4948/ Long 48.0842 | FEN825<br>BI 31131[a] / BI 31132[b] |
| *Dioscorea* species (Ovy valiha) | December 2019 | Lat 12.3731/ Long 49.2448 | FEN828<br>BI 31135[a] / BI 31136[b] |
| *D. alata* | February 2020 | Lat 13.4740/ Long 48.2943 | GHM226<br>BI 31159[a] / BI 31160[b] |
| *D. bako* | February 2020 | Lat 19.98319765/ Long 44.62176857 | MOR12<br>BI 31161[a] / BI 31162[b] |
| *D. buckleyana* | February 2020 | Lat 12.2004/ Long 49.2204 | GHM227 |

| | | | |
|---|---|---|---|
| | | | BI 31143a / BI 31144[b] |
| *D. irodensis* | February 2020 | Lat 12.3419/ Long 49.2447 | FEN860<br>BI 31145[a] / BI 31146b |
| *D. maciba* | February 2020 | Lat 12.2330/ Long 49.1945 | FEN857<br>BI 31141[a] / BI 31142[b] |
| *D. orangeana* | February 2020 | Lat 12.1917/ Long 49.2001 | FEN858<br>BI 31151[a] / BI 31152[b] |
| *D. pteropoda* | February 2020 | Lat 12.2450/ Long 49.2515 | GHM228<br>BI 31155a / BI 31156[b] |
| *D. sambiranensis* subsp. *bardotiae* | February 2020 | Lat 12.2222/ Long 49.1936 | FEN859<br>BI 31149[a] / BI 31150[b] |
| *D. sambiranensis* subsp. *sambiranensis* | February 2020 | Lat 13.2845/ Long 48.4523 | FEN856<br>BI 31147[a] / BI 31148[b] |
| *D. seriflora* | February 2020 | Lat 13.4948/ Long 48.0842 | GHM225<br>BI 31157[a] / BI 31158[b] |
| *Dioscorea* species (Ovy valiha) | February 2020 | Lat 12.3731/ Long 49.2448 | FEN861<br>BI 31153[a] / BI 31154[b] |
| *D. alata* | April - May 2020 | Lat 13.474/ Long 48.2943 | GHM225<br>BI 31248[a] / BI 31249[b] |
| *D. bako* | April - May 2020 | Lat 12.2005/ Long 49.2204 | ND<br>BI 31242[a] / BI 31243[b] |
| *D. buckleyana* | April - May 2020 | Lat 12.342/ Long 49.2446 | GHM228<br>BI 31233[a] / BI 31234[b] |
| *D. irodensis* | April - May 2020 | Lat 12.234/ Long 49.1945 | GHM231<br>BI 31244[a] / BI 31245[b] |
| *D. maciba* | April - May 2020 | Lat 12.1917/ Long 49.2002 | GHM233<br>BI 31240[a] / BI 31241[b] |
| *D. orangeana* | April - May 2020 | Lat 12.245/ Long 49.2515 | GHM229<br>BI 31250[a] / BI 31251[b] |
| *D. pteropoda* | April - May 2020 | Lat 12.2224/ Long 49.1936 | GHM230<br>BI 31238[a] / BI 31239[b] |
| *D. sambiranensis* subsp. *bardotiae* | April - May 2020 | Lat 13.2844/ Long 48.4523 | GHM234<br>BI 31229[a] / BI 31230[b] |
| *D. sambiranensis* subsp. *sambiranensis* | April - May 2020 | Lat 13.4948/ Long 48.0843 | GHM226<br>BI 31252[a] / BI 31253[b] |
| *D. seriflora* | April - May 2020 | Lat 12.3731/ Long 49.2448 | GHM227<br>BI 312486 / BI 31247[b] |
| *Dioscorea* species (Ovy valiha) | April - May 2020 | Lat 13.474/ Long 48.2943 | GHM232<br>BI 31235[a] / BI 31237[b] |
| *D. alata* | June 2020 | Lat 13.474/ Long 48.2943 | GHM242 |

| | | | |
|---|---|---|---|
| | | | BI 31270[a] / BI 31271[b] |
| *D. buckleyana* | June 2020 | Lat 12.342/ Long 49.2446 | GHM238<br>BI 31268[a] / BI 31269[b] |
| *D. irodensis* | June 2020 | Lat 12.234/ Long 49.1945 | GHM236<br>BI 31262[a] / BI 31263[b] |
| *D. maciba* | June 2020 | Lat 12.1917 / Long 49.2002 | FEN864<br>BI 31254[a] / BI 31255[b] |
| *D. orangeana* | June 2020 | Lat 12.245/ Long 49.2515 | GHM237<br>BI 31264[a] / BI 31265[b] |
| *D. pteropoda* | June 2020 | Lat 12.2224/ Long 49.1936 | GHM240<br>BI 31266[a] / BI 31267[b] |
| *D. sambiranensis* subsp. *bardotiae* | June 2020 | Lat 13.2844/ Long 48.4523 | GHM239<br>BI 31272[a] / BI 31273[b] |
| *D. sambiranensis* subsp. *sambiranensis* | June 2020 | Lat 13.4948/ Long 48.0843 | MIR131<br>BI 31258[a] / BI 31259[b] |
| *D. seriflora* | June 2020 | Lat 12.3731/ Long 49.2448 | GHM241<br>BI 31256[a] / BI 31257[b] |
| *Dioscorea* species (Ovy valiha) | June 2020 | Lat 13.474 / Long 48.2943 | GHM235<br>BI 31260[a] / BI 31261[b] |

[a]Parenchyma; [b]Periderm.
